# Elevated methionine induces DNA hypermethylation of transposable elements in *Arabidopsis*

**DOI:** 10.1101/2025.10.30.685479

**Authors:** Yonatan Yerushalmy, Michal Lieberman-Lazarovich, Rachel Amir, Yael Hacham

**Author notes:** Corresponding authors: Rachel Amir, Michal Lieberman-Lazarovich. The author is responsible for the distribution of materials integral to the findings presented in this article in accordance.

## Abstract

Methionine (Met) is a key sulfur-containing amino acid and the precursor of *S*-adenosylmethionine (SAM), the universal methyl donor for DNA and histone methylation. While reduced SAM availability is known to cause DNA hypomethylation, the effects of elevated Met/SAM remain poorly understood. Here, we examined the *Arabidopsis thaliana mto1* mutant, which accumulates Met and SAM due to a mutation in cystathionine γ-synthase, the first committed enzyme of Met biosynthesis. Whole-genome bisulfite sequencing (WGBS) analysis revealed widespread hypermethylation in *mto1*, particularly in non-CG contexts (CHG and CHH), with the strongest changes in pericentromeric heterochromatin. Hypermethylation was concentrated in transposable elements (TEs), especially retrotransposon families such as Gypsy and Copia. Transcriptome profiling showed that TE-genes (TEGs) were broadly downregulated, consistent with reinforced TE silencing due to the hypermethylation. Despite these epigenetic changes, expression of core DNA methyltransferases and demethylases was largely unchanged, suggesting that increased SAM availability enhances enzymatic activity rather than gene expression. In addition, ∼25% of protein-coding genes were differentially expressed in *mto1*, though most changes were not directly linked to promoter- or gene-body methylation. Instead, the strong bias toward hypermethylation of TEs and repression of TEGs suggests that elevated Met/SAM primarily affects gene expression indirectly. Together, these findings demonstrate that an excess of Met/SAM reinforces heterochromatic DNA methylation and transposon silencing, providing new insights into the connection between amino acid metabolism and epigenetic regulation in plants.

## Introduction

Methionine (Met) is a sulfur-containing amino acid (AA) that serves as a fundamental building block for protein synthesis. Its primary metabolic product, *S*-adenosylMet (SAM), is a crucial metabolite in plants, required for essential cellular processes such as the biosynthesis of ethylene, polyamines, vitamins, and phytosiderophores (Amir, 2010). SAM also functions as the main methyl group donor in the synthesis of numerous secondary metabolites, including lipids, alkaloids, and cell wall-related compounds, and is indispensable for the methylation of DNA, RNA, and histone tails (Sauter et al., 2013).

In *Arabidopsis thaliana*, SAM is synthesized by four SAM-synthetase enzymes (Meng et al., 2018). However, previous studies have suggested that SAM synthetase expression is not the primary factor controlling SAM content in plants (Ranocha et al., 2001). Instead, elevated Met levels directly contribute to increased SAM accumulation (Inaba et al., 1994), with 52-80% of Met typically converted into SAM, and the remainder is primarily incorporated into proteins (Giovanelli, 1987; Ranocha et al., 2001). Given the central role of SAM, Met biosynthesis is tightly regulated, in part through a feedback inhibition mechanism whereby SAM represses the transcript of cystathionine γ-synthase (CGS), the first committed and rate-limiting enzyme in the Met biosynthesis pathway (Amir 2010).

Overexpression of *Arabidopsis* CGS (AtCGS) in transgenic tobacco, soybean, potato, and *Arabidopsis* has been shown to substantially increase Met levels (Amir et al., 2019). Furthermore, a modified variant of AtCGS (AtD-CGS), which lacks a 30-amino-acid regulatory N-terminal domain, escapes feedback inhibition by SAM and results in even greater Met accumulation (Hacham et al., 2006). When expressed under a seed-specific promoter, AtD-CGS leads to higher Met content in transgenic soybean, tobacco, and *Arabidopsis* seeds compared to wild-type (WT) controls (Matityahu et al., 2013; Song et al., 2013; Cohen et al., 2014). Unexpectedly, these seeds also accumulate higher levels of soluble metabolites such as sugars, polyols, and amino acids, along with increased starch and total protein, contributing to improved nutritional value.

To investigate the underlying mechanism, seed-specific *AtD-CGS*-expressing *Arabidopsis* lines (SSE plants) were analyzed (Cohen et al., 2014). Transcriptome profiling revealed only a few upregulated genes related to amino acid biosynthesis in seeds, suggesting that the additional metabolites might be transported from vegetative tissues. Supporting this hypothesis, rosette leaves of SSE plants accumulated more Met, sugars and amino acids than control plants with an empty vector (EV) (Girija et al., 2023). Isotope-labeled feeding experiments further demonstrated enhanced flux of aspartate and glutamate from leaves to developing seeds in SSE plants (Girija et al., 2023). Since seeds enriched in Met, proteins, and starch are of nutritional importance, uncovering the mechanism responsible for this metabolic elevation is critical. Further transcriptomic analysis of both leaves and seeds in SSE plants revealed marked changes in the expression of genes associated with DNA and histone methylation. Increased methylation level was subsequently validated using methylation-sensitive enzymes and colorimetric assays (Girija et al., 2023). The results showed that leaves are in a hypermethylation state, and seeds of 21 and 26 days after flowering exhibit equal and lower levels of methylation compared to EV plants, respectively (Girija et al., 2023). These findings formed the basis of our central hypothesis: elevated Met levels lead to increased SAM content, which in turn induces changes in DNA methylation content and therefore can alter the gene expression related to amino acids and sugars.

This study aims to explore the relationship between elevated Met/SAM levels and global DNA methylation patterns, as well as their impact on gene expression. To this end, we investigated the *mto1* mutant, which harbors a mutation in the *AtCGS* gene that reduces feedback regulation by SAM and leads to elevated Met and SAM levels (Inaba et al., 1994). Through integrative methylome and transcriptome analyses, we demonstrate that *mto1* exhibits widespread DNA hypermethylation, particularly in non-CG contexts and heterochromatic regions. Notably, this hypermethylation predominantly affects specific transposable element (TE) superfamilies, especially retrotransposons, and is associated with downregulating TE-associated genes (TEGs). In addition, 25% of all genes significantly changed their expression levels. These results suggest that elevated Met levels contribute to epigenetic silencing of TEs, revealing a novel connection between Met biosynthesis and transcriptional regulation through epigenetic changes.

## Results

### Met content in the *mto1* mutant

The primary aim of this study was to determine whether elevated Met levels are associated with increased DNA methylation. To address this, we utilized the *methionine-overaccumulation 1* (*mto1*) mutant, which is known to accumulate higher levels of Met and SAM (Inaba et al., 1994). To confirm that these metabolite levels remain elevated in our growth conditions, we conducted a Liquid Chromatography–Mass Spectrometry (LC-MS/MS) analysis on leaves from 21-day-old plants. The results demonstrated significant increases in Met, SAM, and *S*-adenosylhomocysteine (SAH) levels in *mto1* compared to WT plants, with fold changes of 11.7, 3.5, and 2.8, respectively (Fig. S1). Consistent with previous findings (Inaba et al., 1994), the *mto1* mutant displayed no marked morphological differences from WT under our experimental conditions.

### *mto1* exhibits increased DNA methylation predominantly in non-CG contexts

To assess the impact of Met/SAM accumulation on global DNA methylation, we initially performed a colorimetric analysis using the MethylFlash Global DNA Methylation (5-mC) ELISA Easy Kit (EpiGentek). The results revealed an ∼8% increase in total 5-methylcytosine (5-mC) levels in *mto1* leaves compared to the WT (Fig. S2).

For a more detailed assessment, we performed whole-genome bisulfite sequencing (WGBS) on genomic DNA from 21-day-old *mto1* and WT leaves. WGBS, a standard method for mapping cytosine methylation in *Arabidopsis*, yielded an average genome-wide mapping rate of 71.4% (Table S1). The conversion rate was calculated based on the chloroplast genome with a conversion rate OF more than 99% for all samples (Table S1). As expected, methylation levels varied by sequence context. In WT, methylation frequencies were 25.82% (mCG), 6.84% (mCHG), and 2.26% (mCHH) (Table S2), consistent with previous reports (Cokus et al., 2008; Feng et al., 2020; Ni et al., 2021). In *mto1*, these values were slightly elevated to 25.95% (mCG), 8.24% (mCHG), and 2.54% (mCHH). Region-specific analysis revealed lower methylation in euchromatin compared with heterochromatin in both WT and *mto1* (Table S2), consistent with earlier studies (Tan et al., 2016). The largest differences between *mto1* and WT were observed in heterochromatin. In euchromatin, *mto1* showed only modest, non-significant increases in methylation across all sequence contexts, as well as in heterochromatic mCG and mCHH. By contrast, heterochromatic CHG methylation increased significantly by 5.3% from 21.7% in WT to 26.7% in *mto1* (Table S2). To further characterize these changes, we identified differentially methylated regions (DMRs) using 100-bp windows and beta regression analysis (padj < 0.05). We focused on DMRs rather than single cytosines because methylation typically acts over contiguous stretches of DNA, such as promoters, gene bodies, and TEs. DMR-based analysis reduces noise from single-site variability, increases statistical robustness, and provides a clearer biological interpretation by linking methylation changes to functional genomic features. DMRs also show whether a region is hyper- or hypomethylated (Catoni et al., 2018). A total of 10,312 DMRs were detected between *mto1* and WT, corresponding to 9,914 unique genomic regions. Most DMRs were found in the mCHG context (4,679), followed by mCHH (3,594), and mCG (2,039). While mCG DMRs were nearly balanced between hyper- and hypomethylation, mCHG and mCHH DMRs were predominantly hypermethylated (Fig. 1A). When normalized per mega-base (Mb), hyper-DMRs outnumbered hypo-DMRs in all contexts and across all chromosomes, with the most pronounced effects observed in non-CG contexts (Fig. 1B).

**Figure 1.**
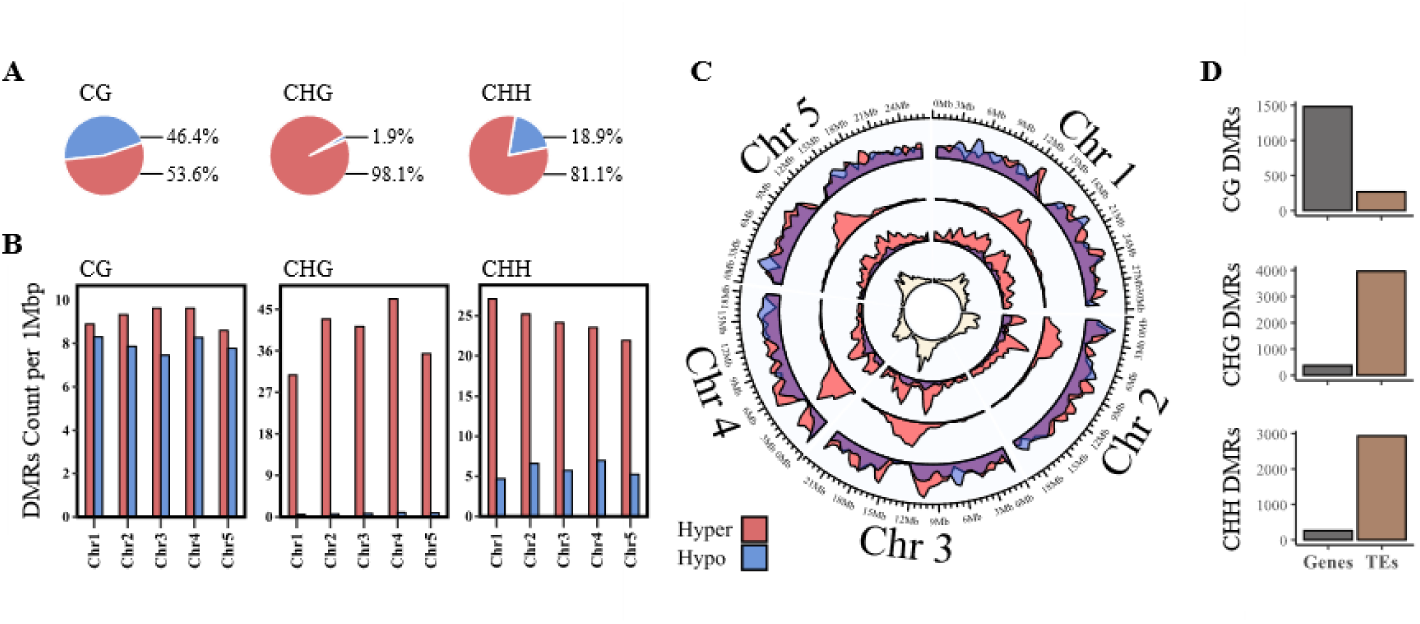
Hyper- and hypo-methylation in the form of DMRs between *mto1* and WT. (A) Pie charts representing the DMRs proportion that were in hyper (red) and hypo (blue) state; (B) The total numbers of DMRs that were in a state of hyper or hypo in each of the five chromosomes in each context of methylation. DMR was counted per 1Mbp; (C) Circus plots visualizing the densities of DMRs across all the chromosomes in CG, CHG, and CHH contexts (from outer to inner circles, respectively). Red indicates hypermethylated regions, blue denotes hypomethylated regions, and purple highlights areas overlapping hyper- and hypomethylated DMRs. In the innermost circle, the density of coding TEs is shown in bright yellow and marks the pericentromeric heterochromatin regions; (D) Count of DMRs in CG, CHG, and CHH contexts, overlapped with coding genes and TEs.

Although the highest levels of DNA methylation were detected in the CG context (Table S2), the differences between *mto1* and WT were relatively modest. This likely reflects the already high baseline methylation of CG sites in WT plants (Fig. 1; Fig. S3) (Stroud et al., 2014). At the genome-wide level, CG-context DMRs (mCG) were broadly distributed but largely excluded from centromeric regions. In contrast, CHG-context DMRs (mCHG) were strongly enriched in these heterochromatic domains (Fig. 1C; Fig. S3). This distribution is consistent with previous reports showing that CHG methylation is typically concentrated in centromeric and pericentromeric regions (Quesneville, 2020; Wang and Baulcombe, 2020), whereas gene-rich, transcriptionally active regions exhibit substantially lower CHG methylation (Fang et al., 2022). Similarly, in *mto1*, CHH-context DMRs were predominantly localized to pericentromeric regions, though their density declined sharply within the centromere and was more diffusely distributed along chromosomal arms (Fig. 1C).

The elevated SAM levels in *mto1* led to changes in specific genomic regions, particularly the centromere and pericentromeric regions (Fig. 1C). To determine whether these locations were expected, we compared our results with published data on Arabidopsis mutants deficient in DNA methylation. We reasoned that regions showing hypermethylation in *mto1* would display hypomethylation in these mutants, due to the consistency of the DNA methylation deposition mechanism. Indeed, analysis of four DNA methylation–deficient mutants (*cmt2, cmt3, met1,* and *ddm1*) (Stroud et al., 2014) revealed opposite methylation patterns, suggesting that the changes observed in *mto1* are both robust and biologically meaningful (Fig. S4).

In *mto1* compared with WT, DMRs corresponding to CHG and CHH methylation were preferentially enriched in TE-dense heterochromatic regions surrounding the centromeres, whereas CG methylation DMRs were largely confined to gene bodies in euchromatin, as expected (Fig. 1D). As shown in Fig. 1C and Fig. S4, CG DMRs comprised both hypo- and hypermethylated regions, a somewhat unexpected result given the elevated Met/SAM level in *mto1*. In contrast, hypermethylation in *mto1* is concentrated primarily at non-CG sites within heterochromatic, TE-rich regions (Fig. S3).

### Elevated Met/SAM levels promote non-CG hypermethylation of retrotransposons

DNA methylation is critical for maintaining genome stability, largely by silencing TEs (Quesneville, 2020; Ito, 2022). To assess TE methylation changes in *mto1*, we generated metaplots for all annotated TEs from the TAIR10 database, including 2 kb upstream and downstream flanking regions (Figs. 2A, B; Fig. S5). As in WT (Law and Jacobsen, 2010), CG methylation was the most abundant context in *mto1*, but its levels were largely unchanged relative to WT (Fig. 2A). In contrast, CHG methylation was markedly elevated in *mto1* (Figs. 2A, B). For CHH sites, the low baseline in WT means that even modest Δ values appear as a substantial relative increase (Fig. 2A). However, when differences relative to WT were calculated, CHH methylation changes were comparable to CG, albeit slightly higher (Fig. 2B). Differential methylation analysis further identified 265, 3,956, and 2,933 TE-associated DMRs in CG, CHG, and CHH contexts, respectively, with most changes occurring in non-CG contexts (Tables S3, S4).

**Figure 2.**
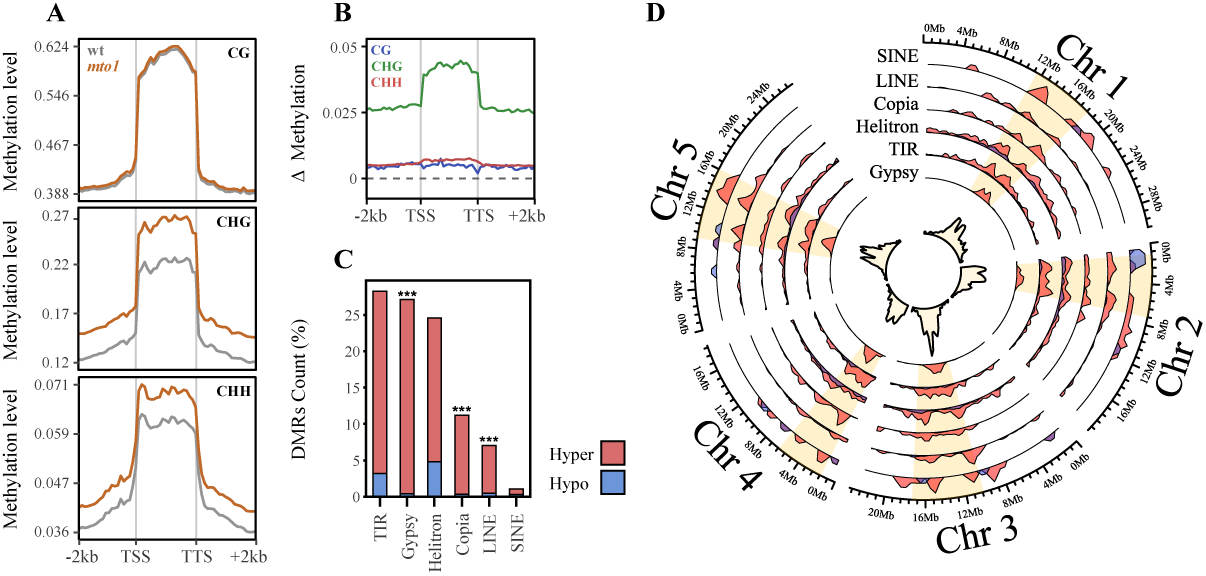
DNA methylation dynamics in transposable elements (TEs). (A) Metaplots of DNA methylation in CG, CHG, and CHH contexts in *mto1* (orange) and WT (gray); (B) DNA methylation difference (Δ, *mto1 vs*. WT) across TEs and their 2 kb upstream/downstream flanking regions. The methylation levels are plotted using mean values across 20 equally sized bins within each region; (C) Distribution of DMRs among major TE superfamilies (Gypsy, Copia, LINE, SINE, TIR, Helitron). The y-axis indicates each superfamily’s proportion of all TE-overlapping DMRs. Enrichment of TE copies (≥1 DMR) was tested against all other TEs using a one-sided Fisher’s exact test. Significance: *p < 0.05; **p < 0.01; ***p < 0.001; (D) Circos plot shows the chromosomal distribution of TE-overlapping DMRs from the six main superfamilies. Red, hypermethylated DMRs; blue, hypomethylated DMRs; purple, overlapping hyper- and hypomethylated regions. The yellow highlights pericentromeric heterochromatin, with TE density shown in the innermost circle. Abbreviations: TSS, transcription start site; TTS, transcription termination site.

Most TE-overlapping DMRs were found in three superfamilies, TIR, Gypsy, and Helitron, together representing 79.9% of all TE-DMRs (28.2%, 27.1%, and 24.6%, respectively; Tables S3-S5). Copia and LINE elements contributed an additional 11.2% and 7.1%, respectively, whereas SINEs and unassigned elements each accounted for <1%. Overall, TE-DMRs were predominantly hypermethylated (90.4%; Table S5). To identify which superfamilies were most strongly affected, we assessed the fraction of elements carrying ≥1 DMR. A one-sided Fisher’s exact test showed significant enrichment of Gypsy, Copia, and LINE elements in *mto1* (Fig. 2C; Tables S3-S5), with DMRs in these superfamilies being hypermethylated. Helitron and TIR elements also displayed a predominance of hypermethylated DMRs, but their enrichment relative to the TE background was not statistically significant (Fig. 2C; Table S5). At the family level, several lineages within the Copia, Gypsy, and LINE superfamilies were specifically enriched for DMRs (Fig. S5). For example, the heat-activated Copia78 element (*ONSEN*) was significantly over-represented in *mto1*.

In *mto1* all the TE superfamilies, the DMRs were more abundant in heterochromatic regions (Fig. 2D, excluding chromosomes 1-2 in SINE). Gypsy elements were predominantly enriched in centromeric and pericentromeric regions, whereas SINE, LINE, Copia, Helitron, and TIR elements were more broadly distributed along heterochromatin edges and chromosome arms, where they may influence gene expression in euchromatic regions (Fig. 2D). Consistent with previous reports (Thao et al., 2014), retrotransposon length was positively correlated with proximity to adjacent heterochromatin regions. In *mto1*, CHG methylation differences relative to WT increased with TE length, with clear hypermethylation in large elements (Fig. S6B), and declined with distance from the centromere (Fig. S6D). By contrast, CG and CHH methylation showed only minor dependence on size or distance (Fig. S6 A-C), consistent with the CHG-dominant effect. Because long TEs are typically associated with heterochromatic regions in *Arabidopsis* (Thao et al., 2014), these size- and distance-dependent trends implicate heterochromatic retrotransposons as the primary targets of hypermethylation in *mto1*.

When TEs were classified by size as long (>4 Kbp) or short (<0.5 Kbp) elements, the differences in methylation between *mto1* and WT were consistently greater for long TEs and their 2-kb flanking regions compared with short elements (Fig. S7A). This effect was observed across all sequence contexts, with the strongest increase in CHG, followed by CHH and CG (Fig. S7A). Within CHG sites, retrotransposons, particularly Gypsy, Copia, and LINE elements, which are generally longer and enriched in heterochromatin, showed the most pronounced hypermethylation (Fig. S7B). In contrast, DNA transposons such as Helitron and TIR, which are typically shorter and located in euchromatin, displayed a weaker effect, even though they are more likely to directly influence nearby gene expression (Fig. S7B).

Together, these findings indicate that elevated Met/SAM levels in *mto1* promote hypermethylation, preferably in retrotransposons, suggesting that the changes are non-random and target specific TE subfamilies.

### Hypermethylated DMRs suppress TEs gene expression in *mto1*

To assess the impact of DMRs on the expression of Gypsy, Copia, and LINE elements, we performed RNA sequencing (RNA-seq) using the same biological samples analyzed by WGBS (Fig. S9). We specifically examined the expression of TE genes (TEGs), which include protein-coding sequences required for transposon mobilization as well as transposon-derived pseudogenes (Feschotte and Pritham, 2007). In total, 1,480 DMRs were mapped to 704 distinct copies of Gypsy, Copia, and LINE TEGs (Fig. 3A; Tables S4, S5), of which 98.4% were hypermethylated in *mto1* relative to WT. Among these TEGs, 117 were differentially expressed (DE-TEGs; *padj* < 0.05), and the vast majority (114; 97.4%) were downregulated (Fig. 3B-C; Table S6). Most DMRs associated with DE-TEGs occurred in non-CG contexts (Fig. S10; Table S6). Notably, 95 of the 117 DE-TEGs (81%) overlapped with DMRs, and 94 out of the 95 DE-TEGs (98.9%) were DMR-hypermethylated in non-CG sites (Tables S6, S7). Consistent with stable TE silencing in WT (Daron and Slotkin, 2017) and the enhanced hypermethylation observed in *mto1*, no evidence of transposition activity was detected in either genotype, based on analysis of unmapped reads across the DE-TEGs (Fig. S7).

**Figure 3.**
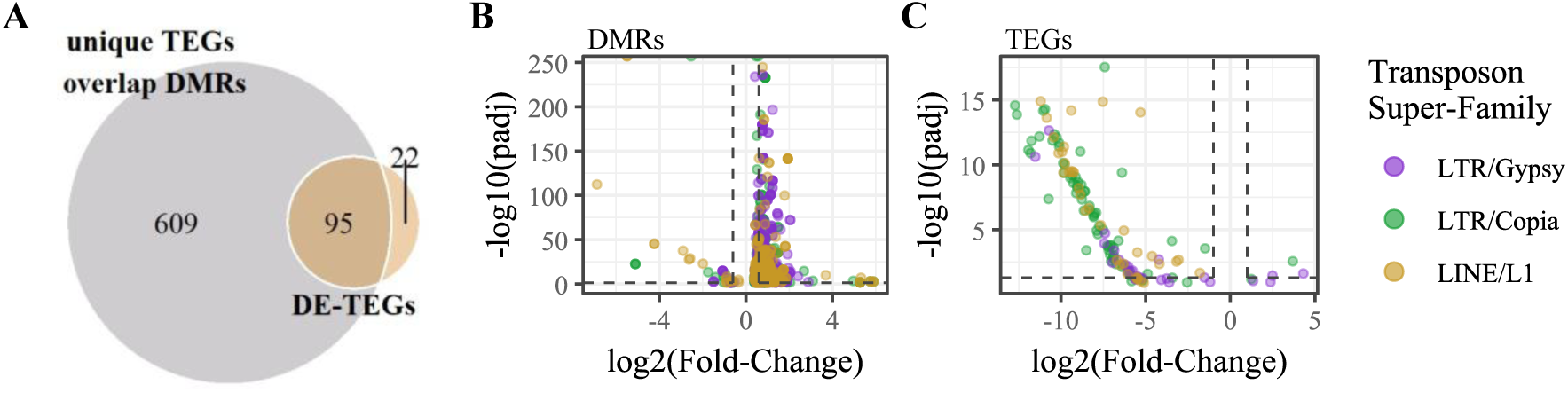
The link between DMRs and transposable element genes (TEGs) expression. (A) Venn diagram showing the overlap between unique TEGs containing at least one DMR and differentially expressed TEGs (DE-TEGs; padj < 0.05). The numbers indicate unique gene IDs in each section of the diagram. (B-C) Volcano plot illustrating the log₂ fold-change and adjusted P-values (*mto1 vs*. WT) for DMRs within TEGs and DE-TEGs. The data represent three retrotransposon superfamilies: Gypsy, Copia, and LINE; TEG lists are based on Tables S6-S7.

Together, these findings strongly suggest that the elevated DNA methylation observed in *mto1*, particularly in non-CG contexts, contributes to the transcriptional repression of Gypsy, Copia, and LINE retrotransposons. This effect is especially pronounced in pericentromeric regions, underscoring the critical role of DNA methylation in restricting TE activity.

### Elevated Met/SAM levels in *mto1* lead to increased DNA methylation in gene bodies and flanking regions

Alterations in DNA methylation were also observed in the euchromatic regions of *mto1* (Fig. 1C), prompting further investigation into the effects of Met/SAM accumulation on gene body methylation and its potential impact on gene expression. We first quantified the number of DMRs and the proportion of hypermethylated DMRs within gene bodies and promoter regions (Tables S8, S9; Fig. S11). The percentage of CG-context DMRs in gene bodies was markedly higher in *mto1* compared to WT (5.73% *vs*. 0.97%). In contrast, promoter regions exhibited a greater proportion of non-CG DMRs than gene bodies, particularly in CHG (2.27%) and CHH (6.86%) contexts. Most CHG-DMRs (89-100%) and CHH-DMRs (78-86%) were hypermethylated, whereas CG-DMRs showed lower levels of hypermethylation (47-73%) (Table 1; Fig. S11).

**Table 1.** The count of DMRs within genes is divided into three DNA-methylation contexts. The percentage of the hypermethylated-DMRs from the total in each context is shown. Genes with at least one DMR in each context were counted (unique genes). The % of hypermethylation for each genic feature is shown near each mC context.

| Genic feature | CG | hyper (%) | CHG | hyper (%) | CHH | hyper (%) | Total | hyper (%) | Unique genes |
| --- | --- | --- | --- | --- | --- | --- | --- | --- | --- |
| Promoters | 900 | 54.3 | 950 | 94 | 2239 | 79.9 | 4089 | 77.5 | 2792 |
| CDS | 1305 | 47.5 | 287 | 96.9 | 114 | 85.9 | 1706 | 58.6 | 1433 |
| Introns | 805 | 48.3 | 195 | 97.4 | 175 | 84.6 | 1175 | 61.9 | 963 |
| 5'UTRs | 26 | 73.1 | 20 | 100 | 14 | 78.6 | 60 | 83.3 | 37 |
| 3'UTRs | 112 | 53.6 | 19 | 89.5 | 42 | 83.3 | 173 | 64.7 | 157 |

To assess broader methylation patterns, we analyzed methylation levels within gene bodies and in 2 kb upstream/downstream flanking regions for all annotated genes (Fig. S12). While CG methylation patterns remained largely unchanged, CHG methylation was elevated in *mto1* across both gene bodies and flanking regions. CHH methylation was also higher in the flanking regions but notably reduced in UTRs and gene bodies of *mto1* compared to WT (Fig. S12). These results indicate that elevated Met/SAM levels in *mto1* induce non-CG hypermethylation in genic and regulatory regions, potentially influencing gene regulation.

### DNA methylation in gene bodies and flanking regions modulates gene expression in *mto1*

To explore the relationship between DNA methylation changes in gene bodies and flanking regions and their impact on gene expression in *mto1*, we analyzed RNA-seq data. Overall, 25% of genes in *mto1* showed significant differential expression compared with WT, with 11% upregulated and 14% downregulated (Fig. S9). To test whether elevated Met/SAM levels influence gene expression through DNA methylation, we examined the overlap between DEGs and DMRs. We found that 9% of DEGs contained promoter-associated DMRs and 7% contained gene-body DMRs (Fig. 4; Table S8-S9). In total, 506 DEGs harbored promoter DMRs, 388 carried gene-body DMRs, and 58 overlapped both categories.

**Figure 4.**
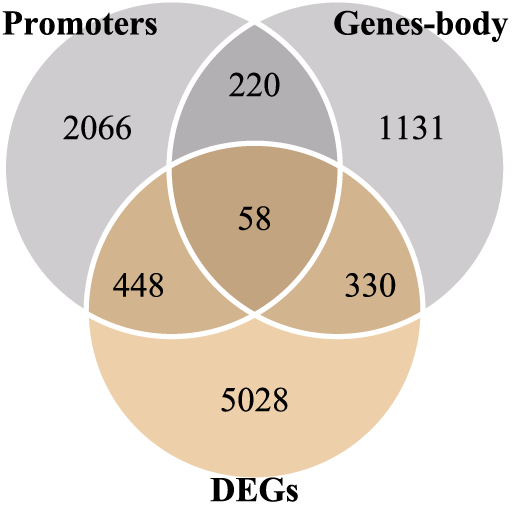
Venn diagram shows the overlap among genes that contain at least one DMR in the promoter or gene-body, and genes that are differentially expressed (DEGs; padj < 0.05). The numbers indicate unique gene IDs in each section of the diagram. The data relies on Tables S8-S9.

To evaluate the impact of DMRs on expression, we analyzed DEGs with at least one DMR (Tables S8-S9). Among these, 18.1% of DMR-containing promoters were associated with altered gene expression. Within genic features, DEGs with DMRs in CDS, introns, 5′ UTRs, and 3′ UTRs were linked to differential expressions at rates of 22.9%, 24.1%, 27%, and 24.8%, respectively (Fig. 5; Tables S8-S9, filtered by unique genes). Overall, 55.4% of DEGs containing DMRs were downregulated (Table S8). CG-DMRs showed balanced hypo- and hypermethylation patterns, while non-CG DMRs were predominantly hypermethylated. Despite this, both up- and down-regulated DEGs were similarly represented (Table S9; Fig. S13), indicating that the methylation in *mto1* might influence expression bidirectionally (Table 2).

**Figure 5.**
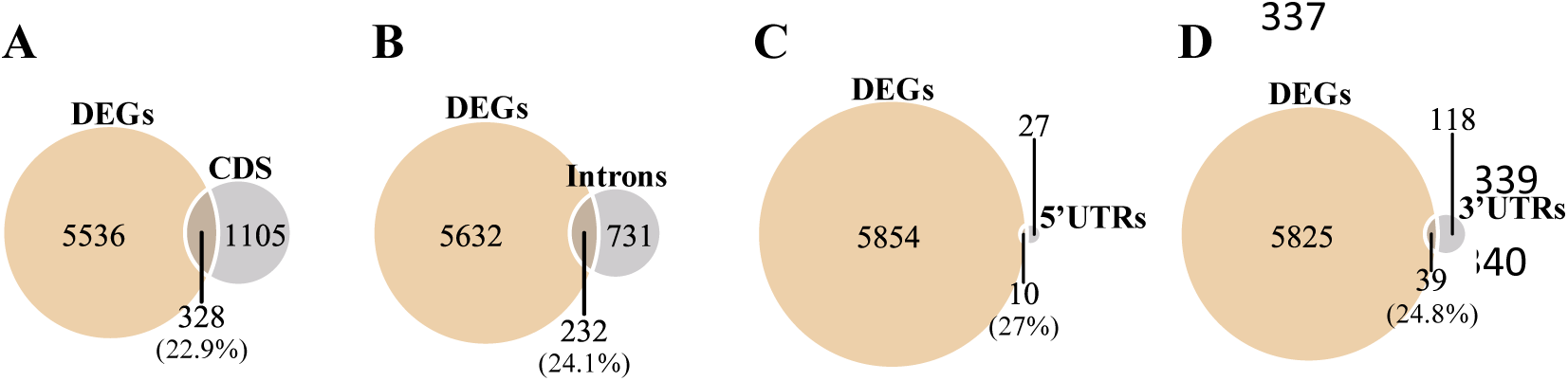
Venn diagram shows the overlap among differentially expressed (DEGs) that contain at least one DMR (orange) and DMRs (gray) in the (A) CDS, (B) introns, (C) 5’-UTRs, (D) 3’-UTRs, (padj < 0.05). The numbers indicate unique gene IDs in each section of the diagram. Percentages represent the proportion of genes with overlapping DMRs and DEG compared to all genes with DMRs in the respective genomic feature. The data relies on Tables S8–S9.

**Table 2.**
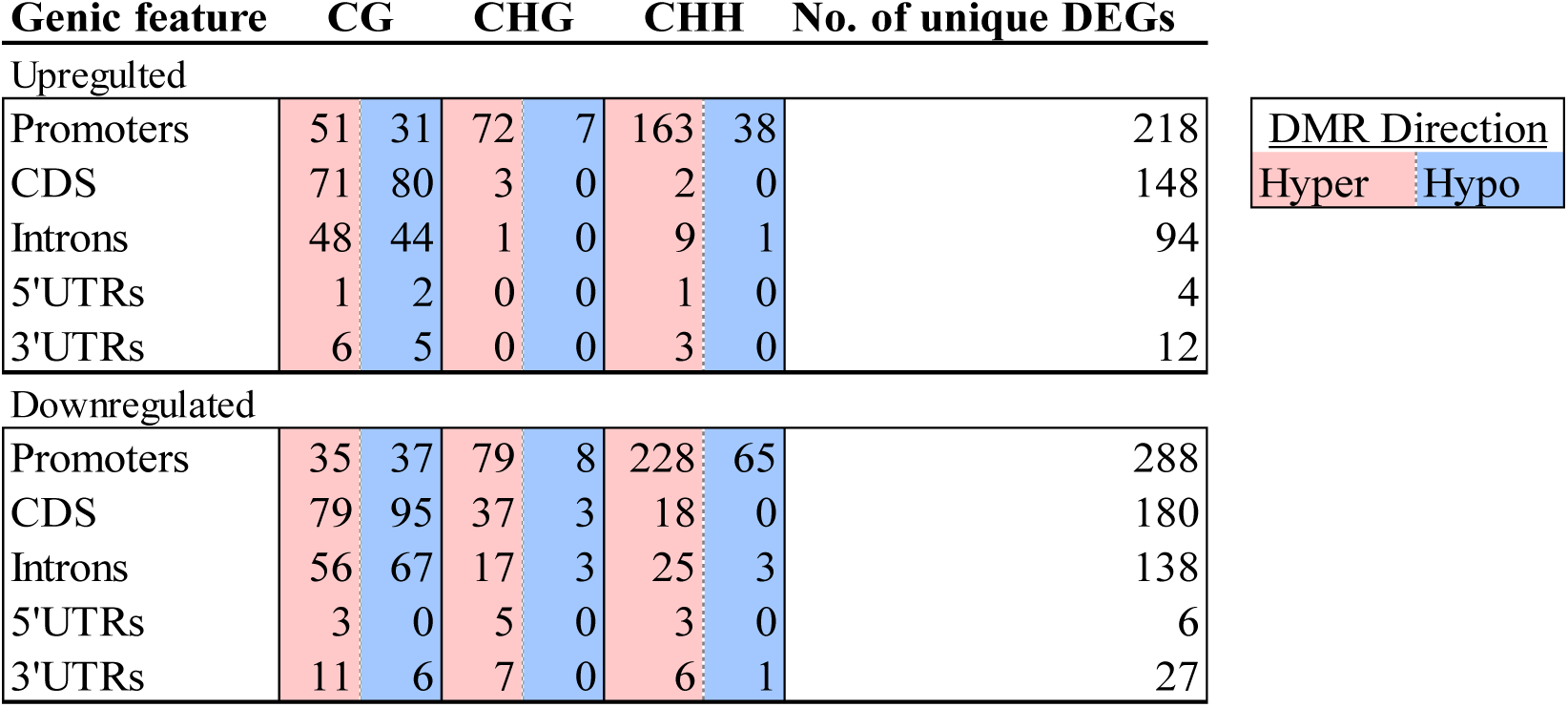
The effect of higher Met/SAM on the hyper-DMRs (red) and hypo-DMRs (blue) in methylation contexts and genic features, overlapping with DEGs (padj < 0.05). The table shows the number of DEGs that overlap with DMRs in each genic feature, separated into upregulated and downregulated categories. There are DEGs with more than one context of DMR; therefore, the number of unique genes was also counted. The table summarizes the data obtained in Tables S8-S9.

Previous studies suggested that CG gene body methylation (GbM) is higher in stable housekeeping genes than in dynamically expressed genes (Williams et al., 2023). However, our analysis showed that differentially expressed stable genes in *mto1* had similar CG GbM levels to dynamic genes (Table S10), implying that GbM may modulate both stable and dynamic gene expression in either direction.

To further assess the impact on gene expression, we correlated average methylation levels across different genic elements, promoters, CDS, introns, 5′ UTR, and 3′ UTR, in each cytosine context with normalized read counts of DEGs, using a negative binomial regression model (Fig. 6). The model incorporated the main effects of methylation and genotype (*mto1* vs. WT), as well as their interaction. In both genotypes, some genes with high promoter methylation showed a negative association with expression, while others displayed a positive correlation (Table 2). Overall, the correlation patterns were similar between *mto1* and WT, and the negative binomial fits captured consistent trends across all contexts (Fig. 6). No significant differences were observed in CG methylation within introns or CDS between genotypes (Fig. 6).

**Figure 6.**
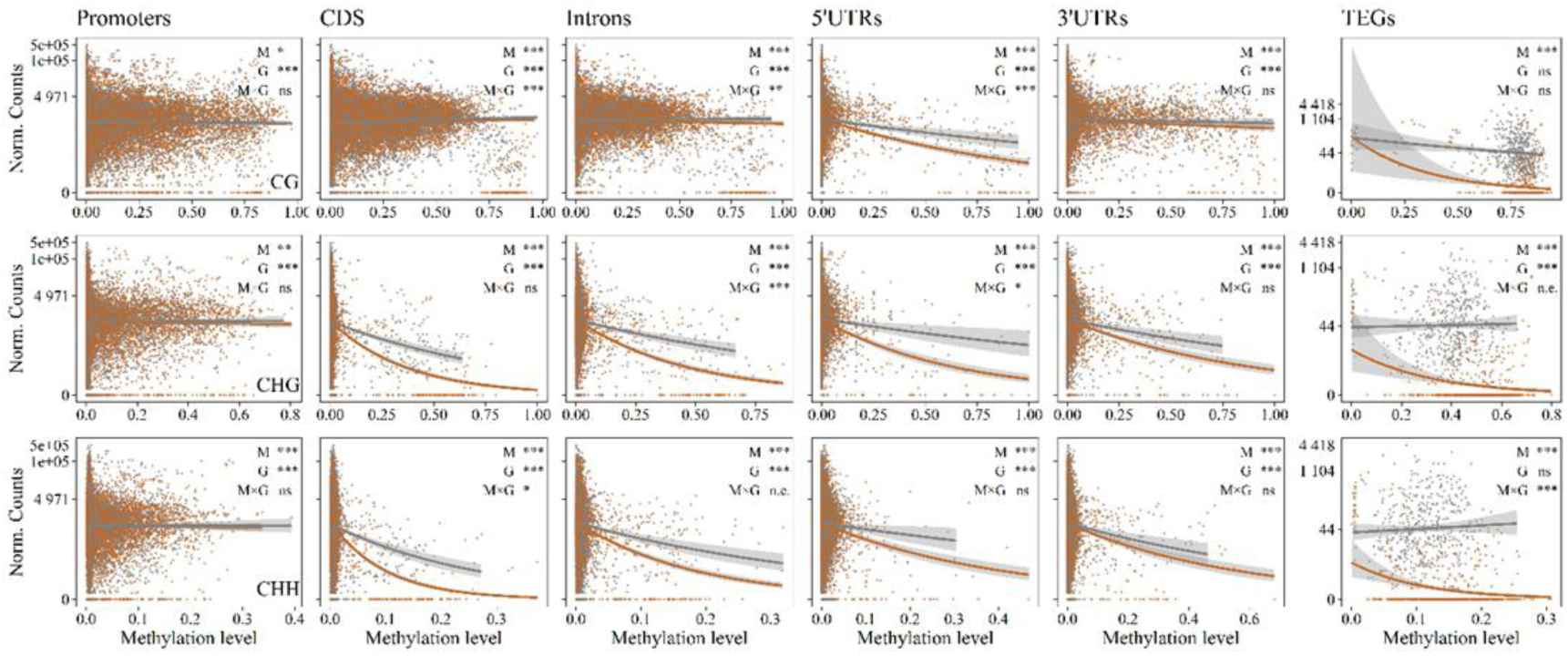
Relationship between average DNA methylation and normalized gene expression count in *mto1* (orange) and wild type (gray). Each point represents a single gene, and smooth curves depict negative-binomial model fits. The Y-axis is a log1p-scaled. Model p-values are provided in Table S11. Bar plots summarizing the dispersion parameter (θ) and deviance-based R² values are shown in Fig. S14. Abbreviations: M, methylation effect; G, genotype effect; M×G, interaction; ns, not significant; n.e., not estimable (due to separation or near-zero counts).

In general, increased non-CG methylation in genic features was associated with reduced expression, particularly in *mto1* (Fig. 6). As a control, we included TEGs in the analysis. As described above (Fig. 3), a strong negative correlation was observed between non-CG methylation and TEG expression, with *mto1* showing consistently higher non-CG methylation and a sharper decline in TEG expression, with several panels not estimable due to near-zero counts (Fig. 6). Despite higher variability in TEG expression (indicated by a lower dispersion parameter, θ), the deviance-based R² was higher than for other gene loci (Fig. S14), suggesting that DNA methylation accounted for a substantial portion of the variation in TEG expression.

To investigate the potential biological impact of promoter and GbM changes in *mto1*, we conducted a Gene Ontology (GO) enrichment analysis of DEGs overlapping with DMRs, focusing on Biological Process terms (Fig. S15; Tables S12-S13). Among the upregulated genes, enriched GO categories included embryo development, RNA modification, leaf senescence, and various metabolic processes. In contrast, the downregulated DEGs were associated with diverse stress responses (including cold, high light, cadmium, and red light), actin filament organization, glycolysis, hypoxia response, and the negative regulation of programmed cell death (Fig. S15).

### Expression levels of DNA methyltransferases and demethylase genes

The *mto1* mutant displays pronounced non-CG hypermethylation relative to the WT, particularly in heterochromatic regions and TEs (Figs. 1, 2; Table S4). In contrast, CG-context DMRs were predominantly enriched within gene bodies (Fig. 1; Tables S2, S7). To evaluate whether this altered methylation landscape could be attributed to increased expression of DNA methyltransferases (MTs), we examined the transcript levels of key DNA MT genes in *Arabidopsis*. Expression analysis of five major MTs revealed no significant differences between *mto1* and WT (Fig. S16; Table S14), suggesting that the hypermethylation phenotype is not due to transcriptional upregulation of these enzymes. We further assessed whether DNA demethylases, which actively remove methylation marks, were affected in *mto1*. Among the four principal demethylase genes examined [DEMETER (DEM) and DEMETER-LIKE 1-3 (DML1-3)], three [DEM, DML1 (also named ROS1), and DML2] showed no substantial change, whereas DML3 expression was significantly downregulated in *mto1* (Fig. S16; Table S14). This reduction in DML3 may contribute to the observed accumulation of non-CG methylation.

## Discussion

This study investigated the potential link between elevated Met and SAM levels and global DNA methylation patterns in *A. thaliana*. While prior studies using mutant lines with reduced SAM levels have demonstrated an association with global DNA hypomethylation (Li et al., 2011; Groth et al., 2016; Meng et al., 2018; Yan et al., 2019), the impact of increased Met/SAM levels on the methylome has remained largely unexplored. The primary objective of this study was to address this knowledge gap. Our results provide compelling evidence that elevated Met/SAM levels lead to widespread changes in both the methylome and transcriptome. Specifically, we present direct evidence that increased SAM availability promotes extensive DNA hypermethylation, primarily in non-CG contexts, with a pronounced enrichment in pericentromeric-heterochromatic regions and TEs (Figs. 1, 2). Like our results, overexpression of Met synthase is also accompanied by a genome-wide global increase in DNA methylation, primarily at the pericentromeric region (González and Vera, 2019). These findings shed light on a previously underappreciated aspect of plant epigenetic regulation via Met/SAM metabolism.

Following the observed global increase in DNA methylation, our analysis revealed a non-random distribution of differentially methylated regions (DMRs) (Fig. 1C). This suggests that certain genomic regions are more susceptible to methylation changes, consistent with prior studies showing that DNA methylation in all contexts is most abundant in heterochromatin and pericentromeric regions (Cokus et al., 2008; Feng et al., 2020; Ni et al., 2021). CG methylation is typically concentrated in centromeric and pericentromeric regions (Cokus et al., 2008; He et al., 2022). In the *mto1* mutant, these regions showed fewer DMR densities in the CG context, and the chromosome arms show balanced DMRs direction, with roughly equal proportions of hyper- and hypo-methylated sites (Fig. 1C). In contrast, mutants such as *met1* and *ddm1* display a dramatic loss of heterochromatic CG methylation (Fig. S4), which also aligns with these findings. Accordingly, the net increase in CG methylation in heterochromatin of *mto1* was negligible (Table S2), likely reflecting the already high baseline of CG methylation in WT.

Unlike CG methylation, non-CG methylation, particularly in the CHG context, was strongly enriched in the pericentromeric regions of *mto1* (Fig. 1C), in agreement with the known localization of CHG methylation in heterochromatin (Cokus et al., 2008; He et al., 2022). These regions are essential for proper chromosome segregation and TE silencing (Feng et al., 2020; Ni et al., 2021). While CHG methylation is concentrated in these regions, CHH methylation is more broadly distributed but also shows high density (Feng et al. 2020; Ni et al. 2021) (Fig. 1C).

Despite the extensive non-CG hypermethylation observed in *mto1*, the transcript levels of the major DNA MTs, CMT2, CMT3, MET1, DRM1, and DRM2, remained unchanged (Fig. S14). CMT2 and CMT3 primarily catalyze CHH and CHG methylation, respectively, with their activity in heterochromatic regions being facilitated by the chromatin remodeler DDM1. In support of this, the *ddm1* mutant shows widespread reductions in CG and CHG methylation and a strong reactivation of TEs (Lee et al., 2023). DRM2 and its homolog DRM3 act in the RNA-directed DNA methylation (RdDM) pathway, guided by small interfering RNAs (siRNAs) to deposit *de novo* methylation at non-CG sites (Fang et al., 2021). Since non-CG methylation in pericentromeric regions is regulated by CMT2, CMT3, and DDM1 (Kenchanmane Raju et al., 2019), we propose that elevated Met/SAM levels in *mto1* may enhance the enzymatic activity of these MTs, rather than increasing their gene expression, thereby contributing to the observed hypermethylation. Additionally, reduced DNA demethylation may play a role, or the involvement of a yet uncharacterized methyltransferase. Among the major demethylases, only DML3 was significantly downregulated in *mto1*. DML3 is crucial for the active removal of non-CG methylation, especially at TEs and genes responsive to stress or senescence, promoting their transcription and developmental roles, such as aging (Schumann et al., 2019). Thus, repression of DML3 may further contribute to the non-CG hypermethylation phenotype in *mto1*.

### The effect of elevated Met/SAM on TE methylation

The elevated non-CG methylation observed in *mto1* heterochromatin occurs primarily at TEs (Fig. 1), which are naturally enriched in heterochromatic regions and are typically heavily methylated (Wang and Baulcombe, 2020; He et al., 2022; Muyle et al., 2022). In contrast to *mto1*, mutants with reduced SAM levels, such as *atms1-1* (methionine synthase) and *mat4* (methionine adenosyltransferase 4), exhibit pronounced hypomethylation in all cytosine contexts, particularly in pericentromeric regions (Meng et al., 2018; Yan et al., 2019). This hypomethylation correlates with strong activation of TEs, highlighting the critical role of DNA methylation in silencing invasive nucleic acids such as TEs, viruses, and transgenes (He et al., 2022).

Beyond their role in genome defense, TEs contribute to epigenetic regulation and can influence genetic and phenotypic variation during development and in response to environmental stresses (Quesneville, 2020; Ramakrishnan et al., 2021). The TE superfamilies most affected by elevated Met/SAM levels in *mto1* include three Class I retrotransposon groups Gypsy, Copia, and LINE, as well as two Class II DNA transposon groups, Helitron and TIR (Papolu et al., 2022) (Table S4; Fig. 2; Fig. S8). Among these, CHG methylation showed a notably higher proportion of hypermethylated DMRs compared to CHH methylation (Table S4; Fig. S6), likely reflecting the more stable maintenance of mCHG through DNA replication, in contrast to the re-establishment required for mCHH in each generation (Lee et al., 2023a). Hypermethylation of TEs in *mto1* was associated with a significant reduction in the expression of TE genes (TEGs) (Table 1; Fig. 3; Fig. 6; Fig. S10), suggesting that increased non-CG methylation enhances TE silencing. It was previously established that no transposition activity is detected in WT plants grown under non-stress conditions (Daron and Slotkin, 2017). Similarly, the hypermethylated *mto1* mutant showed no evidence of transposition, either within the DE-TEG list (Table S7) or specifically for the *ONSEN* element. These findings are consistent with previous studies highlighting the critical role of non-CG methylation in TE repression (Quesneville, 2020; Wang and Baulcombe, 2020; Muyle et al., 2022; Ito, 2022).

All superfamilies that are hypermethylated in *mto1* are predominantly localized in pericentromeric regions, where they contribute to chromatin architecture, chromosome stability, genetic diversity, and regulation of environmental responses (Dupeyron et al., 2019; Protasova et al., 2021). Two feature-level trends reinforce this interpretation. First, in the CHG context, Δ methylation (*mto1*–WT) increased with TE length and declined with distance from the centromere, with the strongest gains observed in large pericentromeric elements (Fig. S6). Second, these elements are primarily heterochromatic retrotransposons, this pattern is consistent with SAM-driven reinforcement of CHG maintenance in heterochromatin. A striking example is the Copia78 (*ONSEN*) retrotransposon, which is normally activated by heat stress and can influence nearby stress-responsive genes through epigenetic mechanisms such as DNA methylation and histone modifications (Deneweth et al., 2022). In *mto1*, *ONSEN* is significantly hypermethylated (Fig. S5), highlighting how elevated Met/SAM levels may further enforce retrotransposon silencing. Copia elements tend to integrate into promoters or introns of stress-related genes, where their activation can either contribute positively to adaptive gene expression or result in deleterious genomic instability, depending on context (Deneweth et al., 2022). Given that TEs comprise a substantial portion of plant genomes, their regulation via methylation is essential for maintaining genomic stability and integrity (Liu and Zhao, 2023). The repression of TEs might reduce the genome plasticity, particularly under stress conditions when TE activation is typically elevated (Ito, 2022).

### High Methionine Content Affects Gene Body Methylation (GbM)

In the *mto1* mutant, non-CG methylation is predominantly enriched in heterochromatic regions and TEs, whereas CG methylation is mainly detected within gene bodies located on the chromosome arms, comprising 5.7% of the total DMRs in *mto1* (Fig. 1D–E). In *A. thaliana*, approximately one-third of all genes exhibit GbM, with ∼20% showing CG methylation specifically in exonic regions (Cokus et al., 2008; He et al., 2022). Although the functional significance of GbM remains under debate (Muyle et al., 2022), it has been proposed to contribute to multiple processes, including the regulation of gene expression, suppression of cryptic transcription, prevention of intron retention, modulation of splicing, and protection from TE insertions (He et al. 2022).

GbM is typically associated with housekeeping genes, which are constitutively expressed at moderate to high levels and are highly conserved across species (He et al., 2022; Muyle et al., 2022; Williams et al., 2023). Notably, GbM genes exhibit reduced transcriptional plasticity across developmental stages and environmental conditions compared to non-methylation genes that show more dynamic expression (Williams et al., 2023). In *mto1*, where Met/SAM levels are elevated, changes in GbM, including both hyper- and hypomethylation, do not consistently correlate with gene expression levels (Tables 2; S8). Similarly, in the *atms1-1* mutant, which exhibits global hypomethylation due to impaired Met synthesis, GbM alterations were not associated with differential gene expression (Yan et al., 2019).

### High Met/SAM levels alter methylation patterns within genic features

Promoter methylation plays a critical role in regulating gene expression. In *A. thaliana*, approximately 5% of genes exhibit methylation in their promoter regions, occurring across all sequence contexts and typically associated with transcriptional repression (Zhang et al., 2006; Kumar and Mohapatra, 2021). However, exceptions to this trend have been reported (He et al., 2022). In the *mto1* mutant, 8.6% of total DMRs are in promoter regions, with 2.27% in the CG context and 6.86% in non-CG contexts. While CG promoter methylation is commonly linked to transcriptional silencing (Niederhuth et al., 2016), in *mto1* these CG DMRs did not show consistent directional effects on gene expression, as both up- and downregulated genes were observed (Table 2, Fig. 6).

Non-CG methylation within promoters is also often associated with transcriptional repression (Niederhuth et al., 2016; Wang and Baulcombe, 2020). Like CG methylation, CHG hypermethylation induced by elevated Met/SAM levels did not correlate with expression changes (Table 2). In contrast, DMRs of CHH were more than twice as frequent and predominantly hypermethylated, coinciding with a modest increase in the number of downregulated genes (Table 2; Table S8). Elevated CHH methylation at promoters has also been observed under abiotic and hormonal stresses, such as in strawberries (Wang and Baulcombe, 2020; López et al., 2022), suggesting potential regulatory flexibility. The relationship between promoter methylation and gene expression is species dependent. In rice, CG and CHG methylation typically represses gene expression, while CHH methylation correlates positively (Wang and Baulcombe, 2020). In contrast, no clear association was found in apples (*Malus domestica*) (Xu et al., 2018).

Within gene bodies, most DMRs were in CDS and introns. In *mto1*, CG DMRs included both hypo- and hypermethylated sites, showing no consistent association with expression. By contrast, most non-CG DMRs were hypermethylated and inversely correlated with expression, with stronger effects in CHG than CHH (Table 2; Fig. 6). Across 5′UTRs, CDS, and introns, methylation– expression relationships in *mto1* were lower, with steeper slopes than in WT. Thus, non-CG hypermethylation in the mutant generally coincides with reduced transcription. TEGs were particularly affected, consistent with reinforced silencing (Figs. 6, S14). This reduction in expression may limit transcriptional flexibility, potentially decreasing the ability of *mto1* to tolerate stress. Supporting this view, GO enrichment analysis revealed that many downregulated genes in *mto1* are involved in stress responses (Fig. 6). Similarly, overexpression of Met synthase, which elevates DNA methylation, has been shown to repress plant immunity and increase susceptibility to *Pseudomonas syringae* (González and Vera, 2019). Further studies will be required to validate the proposed link between hypermethylation, reduced expression plasticity, and heightened environmental sensitivity.

## Conclusion

This study establishes a link between elevated Met/SAM levels and increased DNA methylation, particularly in the non-CG context. The DMRs were not randomly distributed but were predominantly enriched in centromeric and pericentromeric heterochromatin, which is rich in TEs. In *mto1*, the number of methylated Class I retrotransposons was significantly higher compared to WT, coinciding with a substantial reduction in the expression of TEGs, including members of the ATHILA family and the heat-responsive element ONSEN, both implicated in epigenetic regulation and stress responses (Deneweth et al., 2022). The inverse relationship between increased methylation and reduced TEG expression, supported by GO enrichment analysis, suggests that TE silencing may limit the plant’s ability to adapt to environmental stress by altering chromatin dynamics. Nevertheless, this hypothesis requires further validation through targeted stress assays in the *mto1* mutant.

One potential explanation for the preferential hypermethylation of heterochromatin is related to Met metabolism. Elevated Met is known to impair plant growth and development (Hacham et al., 2002; Xiang et al., 2022). To mitigate toxicity, plants catabolize Met into volatile compounds or divert its intermediates into the biosynthesis of secondary metabolites such as glucosinolates, polyamines, *S*-methylMet, and isoleucine (Hacham et al., 2002; Cohen et al., 2014; Hacham et al., 2023). Nonetheless, the precise mechanisms of Met toxicity remain elusive. Recent findings show that increased Met biosynthesis impairs the plant’s ability to cope with oxidative stress (Hacham et al., 2024). The current study suggests a mechanism for managing the excess of Met/SAM by directing methyl groups toward heterochromatin regions that are less essential for daily growth and development than euchromatin. This redistribution might represent an adaptive buffering strategy to protect the epigenome from high SAM content and methylation overload. However, the widespread transcriptional changes observed in *mto1* indicate that additional, yet unidentified, mechanisms may be involved. These changes could reduce *mto1* fitness relative to WT, even under standard growth conditions.

In summary, our findings highlight the influence of elevated Met/SAM levels on epigenetic regulation in plants and suggest a novel connection between primary metabolism and chromatin dynamics. Future research should aim to dissect the mechanistic links between SAM abundance, MTs activity, chromatin remodeling, and plant stress resilience.

## Materials and Methods

### Plant genotypes and growth conditions

The Arabidopsis (*A. thaliana*) mutant lines of *mto1* (CS196) were obtained from ABRC, Ohio State University, USA. *mto1* and WT (*Col-0*) plants were grown in 0.5 Murashige and Skoog media (DuShefa, Haarlem, The Netherlands) supplemented with 1% sucrose under a 16-h light/ 8-h dark cycle in a growth chamber (22 °C ± 2 °C). The plants were subsequently transferred to soil and placed in a growth chamber under the same light conditions at 22 °C ± 2 °C.

### Metabolite detection

21-day-old Arabidopsis leaves were frozen in liquid nitrogen and lyophilized. The lyophilized tissues were ground to a fine powder in liquid nitrogen and kept at −80°C until use. Met, SAM, and SAH were extracted from 50 mg of dry powder. The samples were analyzed by injecting 5 µL of the extracted solutions (100 mg/mL of aqueous solution with 0.01N HCl) into an Ultra-High Performance Liquid Chromatography (UHPLC) connected to a photodiode array detector (Dionex Ultimate 3000), with a reverse-phase column (Phenomenex RP-18, 100-3.0 mm, 2.5 μm). The mobile phase consisted of (A) DDW with 0.1% formic acid and (B) acetonitrile containing 0.1% formic acid. The gradient started with 100% A for 1 min, then increased to 30% B in 7 min, then to 90% B in 2 min, and kept at 90% B for another 3 min. Phase A was returned to 100% A in 1 minute, and the column was allowed to equilibrate at 100% A for 2 min before the next injection. The flow rate was 0.5 mL/min. LC-MS/MS analysis was performed with a Heated Electrospray ionization (HESI-II) source connected to a Q Exactive™ Plus Hybrid Quadrupole-Orbitrap™ Mass Spectrometer Thermo Scientific ™. ESI capillary voltage was set to 3500 V, capillary temperature to 300 °C, gas temperature to 350 °C, and gas flow to 10 mL/min. The mass spectra (m/z 100–1500) were acquired in positive-ion mode.

### Whole-Genome Bisulfite Sequencing (WGBS)

Genomic DNA from leaves of three biological replicates of *mto1* and two of WT was extracted using a DNeasy Plant Mini Kit (Qiagen). We generated global DNA methylation profiles at single-nucleotide resolution using the Illumina Hi-Seq 2500 platform (BGI, Tech Solutions, Hong Kong). DNA quality and contamination were monitored by running samples on 1% agarose gels. From each sample, one ug of genomic DNA was sent to BGI (BGI, Shenzhen, China) for further preparation and analysis.

Genomic DNA was prepared for library construction by fragmenting to an average size of 200–350 bp, followed by blunt-ending, 3′-dA addition, and ligation of methylated adaptors. The libraries underwent bisulfite conversion using the EZ DNA Methylation-Gold Kit (ZYMO), PCR amplification, quality control, circularization, DNA nanoball (DNB) formation, and sequencing on the DNBSEQ platform (DNBSEQ Technology). Raw reads were processed using SOAPnuke (Chen et al., 2018) with the following filtering parameters: removal of adaptor sequences (≥ 25% match, ≤ 2 mismatches), reads < 150 bp, reads containing ≥ 0.1% ambiguous bases (N), reads with polyX stretches > 50 bp, or reads with ≥ 40% of bases having Phred scores < 20. After filtering, only high-quality readings (Phred+33 encoding) were retained for downstream analysis.

### RNA sequencing (RNAseq)

Total RNA was isolated from the leaves of three *mto1* and WT plants using the Spectrum Plant Total RNA kit (Sigma) according to the manufacturer’s protocol. The quality and integrity of the RNA were checked by running samples on 1% agarose gels and using a standard NanoDrop.

RNA-seq libraries were prepared at the Crown Genomics Institute of the Nancy and Stephen Grand Israel National Center for Personalized Medicine, Weizmann Institute of Science using the INCPM-mRNA-seq. Briefly, the polyA fraction (mRNA) was purified from 500 ng of total input RNA, followed by fragmentation and the generation of double-stranded cDNA. After Agencourt Ampure XP beads cleanup (Beckman Coulter), end repair, A base addition, adapter ligation, and PCR amplification were performed. Libraries were quantified by Qubit (Thermo Fisher Scientific) and TapeStation (Agilent). Sequencing was done on a Novaseq 6000 instrument (Illumina) using an S1 100 cycles kit, allocating 40-50M reads per sample (paired-end sequencing).

### Data Analysis

All analyses used the *A. thaliana* TAIR10 genome assembly and annotations. TEs were grouped by transposition mechanism following an Arabidopsis-focused framework (Quesneville, 2020): Gypsy, Copia, LINE, and Helitron were treated as separate groups; all terminal inverted repeat (TIR) superfamilies, such as En-Spm, MuDR, Harbinger, hAT, Pogo, Mariner, and Tc1 were grouped into a single ‘TIR’ category; SINE elements annotated as Rath (RathE*_cons) were included within the SINE order; elements annotated as ‘Unassigned’ were excluded from downstream analyses. TEGs were defined as gene models annotated with type ‘transposable element gene’ in TAIR10. Each TEG was linked to its source TE via the GFF3 ‘Derives_from’ attribute. To examine the transition activity, we used new non-reference TE insertions called from unmapped WGBS reads using epiTEome (default settings; ≥5 supporting split reads; evidence from both TE ends) against the TAIR10 genome and TE annotations (Daron and Slotkin, 2017). Methylome WGBS reads were aligned to the TAIR10 genome using Bismark software (Krueger and Andrews, 2011). Per-cytosine methylation levels were derived from Bismark cytosine calls as 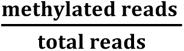 at each covered cytosine. Average methylation (%) was the mean of these per-C values across all covered cytosines within a region per sample and summarized as mean ± SD across replicates. For meta-plots, we oriented all loci as 5′→3′, split each protein-coding gene (or TE) and the ±2 kb flanks regions into 20 length-normalized bins (60 total), computed per-bin mean methylation level for CG/CHG/CHH contexts using sites with ≥6× coverage, then averaged bin-wise across all loci; genic-features metaplots used 10 bins with ≥15 cytosines per bin, otherwise identical. DMRs were calculated using the *DMRcaller* R package (Catoni et al., 2018) in 100-bp bins, applying minimum methylation difference thresholds of 0.4, 0.2, and 0.1 for the CG, CHG, and CHH contexts, respectively, with a ≥6× coverage and ≥4 cytosines per site. Enrichment tests (one-tailed) were performed by comparing the DMR counts in each group to those in its corresponding larger group, with p-values and scores calculated using PlanTEnrichment method (Karakülah and Suner, 2017). Heterochromatin regions were defined as the pericentromeric intervals as previously define (Bi et al., 2017), on TAIR10: Chr1 ∼11.5-17.7 Mb; Chr2 ∼1.1-7.2 Mb; Chr3 ∼10.3-17.3 Mb; Chr4 ∼1.5-6.3 Mb; Chr5 ∼9.0-16.0 Mb; all other sequence was considered euchromatin. For RNAseq data, clean reads were mapped to the Arabidopsis genome using RSEM (Li and Dewey, 2011) to generate normalized expression values for each gene. They differentially expressed genes (DEGs, padj < 0.05) were identified with DESeq2 (Love et al., 2014). To ensure reliable quantification, only genes with normalized read counts >10 and TEGs with >20 were included in the analysis. Representative IGV snapshots of TE-associated DMRs alongside normalized RNA-seq read coverage are shown in Fig. S17, with corresponding values provided in Table S7. For both DMR and DEG analyses, statistical significance was adjusted for multiple testing using the Benjamini–Hochberg correction (Love et al., 2014; Catoni et al., 2018). Negative binomial regression was used to model normalized gene expression read counts as a function of average methylation level across gene bodies and promoter regions while accounting for genotype differences (*mto1 vs*. WT). Gene Ontology (GO) enrichment analysis was also performed using the *topGO* R package with the ‘*weight01*’ algorithm. All data visualizations and statistical analyses were conducted using R (R Core Team, 2018). The R scripts used for the various analyses in this study are available on GitHub at https://github.com/Yo-yerush.

## Data availability

All high-throughput sequencing data from this study are available in the NCBI BioProject database under the accession number: PRJNA1206675. Additional supporting data can be requested from the corresponding authors. The Arabidopsis reference genome (TAIR10), including gene and transposable element annotations, was obtained from The Arabidopsis Information Resource (TAIR; www.arabidopsis.org).

## Acknowledgement

This work was supported by a grant from the Israel Science Foundation (ISF grant no. 1857/20).

## Author contributions

YH, ML, and RA: experimental design. YY and YH: conducted experiments. YY and YH: data analysis. RA: preparing the first draft. YY, YH, ML, RA: improved the manuscript preparation. All the authors read, revised, and approved of the manuscript.

## Supplemental Data

### Supplemental Tables

**Table S1.** Mapping and quality statistics for paired-end WGBS analysis

**Table S2.** Average methylation rate (%) of WT (Col-0) and *mto1*

**Table S3.** DMRs of 100bp overlapping various TEs across all sequence contexts

**Table S4**. The number of DMRs that overlap with various TE superfamilies

**Table S5.** TE superfamily enrichment and composition of TE-overlapping DMRs in *mto1 vs*. WT. DMRs of 100bp overlapping TEGs across all sequence contexts

**Table S6.** Differentially expressed TEGs classified to Gypsy, Copia, and LINE retrotransposon superfamilies

**Table S7.** DMRs of 100bp overlapping coding sequences (CDS), introns, untranslated regions (UTRs), and promoter regions across all sequence contexts

**Table S8.** Differentially expressed genes (DEGs, padj < 0.05) that overlap with at least one DMR in gene-body or promoter regions

**Table S9.** Number of DMRs in various methylation contexts and gene features overlapping Stable-GbM genes that found in DEGs

**Table S10.** P-values for the correlations between DNA-methylation level and genotype (*mto1 vs*. WT)

**Table S11.** Gene Ontology (GO) enrichment analysis for upregulated DEGs overlapping DMRs

**Table S12.** Gene Ontology (GO) enrichment analysis for downregulated DEGs overlapping DMRs

**Table S13.** Differential expression of DNA methyltransferase and DNA demthylase genes in *mto1* relative to WT

### Supplemental Figures

**Figure S1.** The level of Met, SAM and SAH in *mto1* and WT as measured by using LC-MS/MS

**Figure S2.** Change in the percentage of DNA methylation in leaves of *mto1* compared to WT plants, as obtained with MethylFlash™ Global DNA Methylation (5-mC) ELISA Easy Kit

**Figure S3.** Genome-wide methylation levels for *mto1* and WT, shown in 150-kbp sliding windows across the five chromosomes

**Figure S4.** Genome-wide distribution plots depicting methylation differences between *mto1*, various methyltransferase mutants

**Figure S5.** Differences (<u>Δ</u>, *mto1 vs* WT) in TEs

**Figure S6.** Metaplots of DNA methylation difference (Δ, *mto1 vs*. WT) in CG, CHG, and CHH contexts detected at TEs and their flanking 2 kb upstream/downstream regions

**Figure S7.** Enrichment analysis comparing the proportion of TEs with DMRs in each family to the total TEs with DMRs in their respective TE group

**Figure S8.** Pie chart depicting the distribution of DMRs across TE superfamilies

**Figure S9.** Overview of the RNAseq-based transcriptome analysis (*mto1* vs. WT)

**Figure S10.** Volcano plot displaying the fold-change (*mto1* vs. WT) DMRs within TEGs in the three contexts, from the retrotransposon superfamilies

**Figure S11.** Bar plot showing DMRs overlapping with promoters and genic features in CG, CHG, and CHH methylation contexts in *mto1 vs* wild type (WT)

**Figure S12.** Metaplots of mean DNA methylation level calculated from the WGBS data for WT and *mto1* in CG, CHG, and CHH

**Figure S13.** Metaplots of DNA methylation in different sequence contexts, overlapped with genes that belong to DEGs in *mto1* and WT

**Figure S14.** Bar plot of the dispersion parameter (θ) from the negative binomial model fitted to each genomic feature in different DNA methylation contexts

**Figure S15.** Gene Ontology enrichment analysis of DEGs associated with biological process terms that overlap with DMRs in their promoters and gene bodies

**Figure S16.** Bar plot showing the differential expression of five methyltransferase genes and four demethylases in *mto1* relative to wild-type

**Figure S17.** IGV snapshot of DMRs results (*mto1 vs*. WT) and RNA-seq coverage for a representative TE gene.

